# ModA phasevarions regulate adherence of non-typeable *Haemophilus influenzae* to the host airway in a tissue-specific manner

**DOI:** 10.1101/2022.04.13.488267

**Authors:** Preeti Garai, John M. Atack, Brandon M. Wills, Michael P. Jennings, Lauren O. Bakaletz, Kenneth L. Brockman

## Abstract

Adherence of non-typeable *Haemophilus influenzae* (NTHi) to the host airway is an essential initial step for asymptomatic colonization of the nasopharynx, as well as development of disease. NTHi relies on strict regulation of multiple adhesins for its pathogenesis. The ModA phasevarion is a bacterial regulatory system important for virulence of NTHi. However, the role of the ModA phasevarion in adherence of NTHi to the host airway is not understood well. This study addressed the role of the ModA phasevarion in the regulation of adherence of NTHi to multiple substrates of the host airway. Assessment of adherence of the *modA* variants of four clinical isolates of NTHi showed that ModA phasevarions regulated adherence of NTHi to mucus, middle ear epithelial cells, and vitronectin in a substrate-specific manner. The adhesins Protein E and P4 were found to contribute to the ModA-regulated adherence of NTHi to distinct substrates. A better understanding of such tissue-specific regulation of NTHi adherence by the ModA phasevarion will allow identification of virulent NTHi populations at the site of disease within the host airway and facilitate more directed development of vaccines or therapeutics.

## Introduction

Non-typeable *Haemophilus influenzae* (NTHi) is a host-adapted mucosal pathogen that colonizes the human nasopharynx asymptomatically from early childhood (1–3). As a predominant pathogen of the human respiratory tract, NTHi causes infections at multiple sites within the airway. These include otitis media (OM), an inflammatory disease of the middle ear (4), and exacerbations in the lungs of patients with chronic obstructive pulmonary disease (COPD) and cystic fibrosis (CF) (5, 6). Additionally, NTHi causes bronchitis, sinusitis and community acquired pneumonia (7–9). Approximately 10% of the human population (over 700 million people) is affected by OM, which can result in hearing loss and thereby affect learning. The complications associated with OM lead to the death of more than 20,000 people per year globally, with the highest morbidity and mortality in children under the age of 5 (10). On the other hand, COPD is the third leading cause of deaths worldwide (11, 12). There are currently no vaccines available against NTHi.

Adherence of NTHi to the airway epithelium is the primary step in colonization of the human nasopharynx as well as in the development of disease (13–18). For instance, ascension of NTHi from the nasopharynx to the middle ear during the development of OM depends on the adherence of NTHi to mucus present within the Eustachian tube (19). Therefore, the regulation of adherence of NTHi to different host substrates found within the respiratory tract is essential during both commensalism and disease.

NTHi expresses multiple surface associated adhesins that specifically bind to cognate host receptor proteins, extracellular matrix components (ECM) or mucus in order to adhere to different parts of the host airway (19–23). Because NTHi lack a capsule, adhesins are more accessible on the surface of these bacterial cells than their encapsulated counterparts. This accessibility also makes these surface-exposed adhesins good vaccine candidates for protection against NTHi infections (24–26).

NTHi, like several other host-adapted mucosal pathogens, encodes a DNA methyl transferase, ModA that is phase-variable (27–31). Phase variation is the random and reversible switching of expression of a protein. In the case of the encoding *modA* gene, this is due to simple DNA sequence repeats (SSRs) within its open-reading frame. Variation in length of this SSR tract results in the expression (status ‘ON’) or the absence (status ‘OFF’) of the ModA protein. The ModA protein can bind to and methylate multiple sites within the bacterial genome, resulting in genome wide methylation differences commensurate with methyltransferase expression. This differentially regulates the expression of multiple genes via epigenetic mechanisms. Therefore, the status of ModA can affect the expression of a distinct set of genes forming a unique **phase vari**able regul**on** or phasevarion (31). Of the 22 *modA* allelic variants reported for NTHi so far, *modA2* is the most prevalent amongst multiple clinical isolate collections (27, 32, 33). Analysis of phase-variable *modA* allele distribution across multiple NTHi strain collections revealed that two-thirds of the OM isolates encode just one of the following 5 alleles - *modA2, modA4, modA5, modA9*, and *modA10* (33), whereas the majority of the COPD isolates contained a *modA2, modA4*, or *modA5* allele (32). Each ModA phasevarion regulates a unique set of genes, many of which encode adhesins or potential vaccine antigens (32, 33).

The ModA2 phasevarion has been shown to regulate multiple disease-related phenotypes including resistance to oxidative stress (34), biofilm formation (35) and virulence in the chinchilla model of experimental OM (36, 37). The ModA4 and ModA5 phasevarions are known to play distinct roles in the pathogenesis of NTHi such as evasion from opsonization and susceptibility to antibiotics (33). However, the role of the ModA phasevarions in adherence of NTHi to the host airway has not been investigated. This study addressed the roles of the ModA2, ModA4, ModA5, and ModA9 phasevarions in the adherence of NTHi to various cellular and non-cellular substrates encountered by NTHi within the airway. Distinct phasevarions regulated the adherence of NTHi to specific host airway substrates.

## Results

### ModA2 regulates adherence of NTHi to mucus

The NTHi clinical strains 723, C486, 477 and 1209 contain the unique *modA* alleles *modA2, modA4, modA5* and *modA9*, respectively (33), and were therefore selected for this study to represent each of these phasevarions. These strains were genetically modified to permanently lock the expression of ModA in the OFF or ON status, resulting in *modA* locked OFF and *modA* locked ON variants of each strain (Table 1). The use of these locked variants, hereafter referred to as *modA2* OFF and *modA2* ON variants, allowed the assessment of the direct effects of each *modA* status on the adherence of NTHi.

**Table 1:** Bacterial strainsTable.

| Strain | Reference |
| --- | --- |
| NTHi 723 *modA2* locked OFF | (34) |
| NTHi 723 *modA2* locked ON | (34) |
| NTHi C486 *modA4* locked OFF | This study |
| NTHi C486 *modA4* locked ON | This study |
| NTHi 477 *modA5* locked OFF | This study |
| NTHi 477 *modA5* locked ON | This study |
| NTHi 1209 *modA9* locked OFF | This study |
| NTHi 1209 *modA9* locked ON | This study |
| NTHi 723 $\Delta ompE$ *modA2* locked OFF | This study |
| NTHi 723 $\Delta ompE$ *modA2* locked ON | This study |
| NTHi 723 $\Delta hel$ *modA2* locked OFF | This study |
| NTHi 723 $\Delta hel$ *modA2* locked ON | This study |
| NTHi 723 $\Delta hap$ *modA2* locked OFF | This study |
| NTHi 723 $\Delta hap$ *modA2* locked ON | This study |

Mucus coats the surface of the airway epithelium, and as such serves as an initial substrate for adherence by NTHi that can further facilitate movement of the pathogen to different sites within the airway (19, 20, 22, 23, 38). Therefore, adherence to mucus is crucial for NTHi colonization and pathogenesis within the respiratory tract. The respiratory tract epithelium is composed of a variety of cells, which includes goblet cells that produce mucus, and ciliated cells that propel the mucus through the airway (39). Normal human primary bronchial-tracheal epithelial cells (nhPBTEs) were grown at an air-liquid interface to induce formation of differentiated polarized cells that mimic the pseudostratified epithelium of the respiratory tract (40). Mucus produced by these polarized nhPBTEs was then collected and used to assess the adherence of each *modA* variant. Bacteria were incubated in mucus-coated wells for 1 hour and the adherent bacteria were enumerated. The *modA2* ON variant of strain 723 adhered to mucus significantly more than the *modA2* OFF variant (Fig. 1, p<0.001). However, there was no significant difference between the *modA* variants of strains C486, 477 or 1209. Interestingly, strain 477 adhered the least and strain 1209 had the greatest overall adherence to mucus irrespective of *modA* status (Fig. 1). Thus, ModA2 regulates adherence of NTHi to mucus, indicating an important role of the ModA2 phasevarion in adherence of the pathogen to the airway epithelium.

**Fig. 1.**
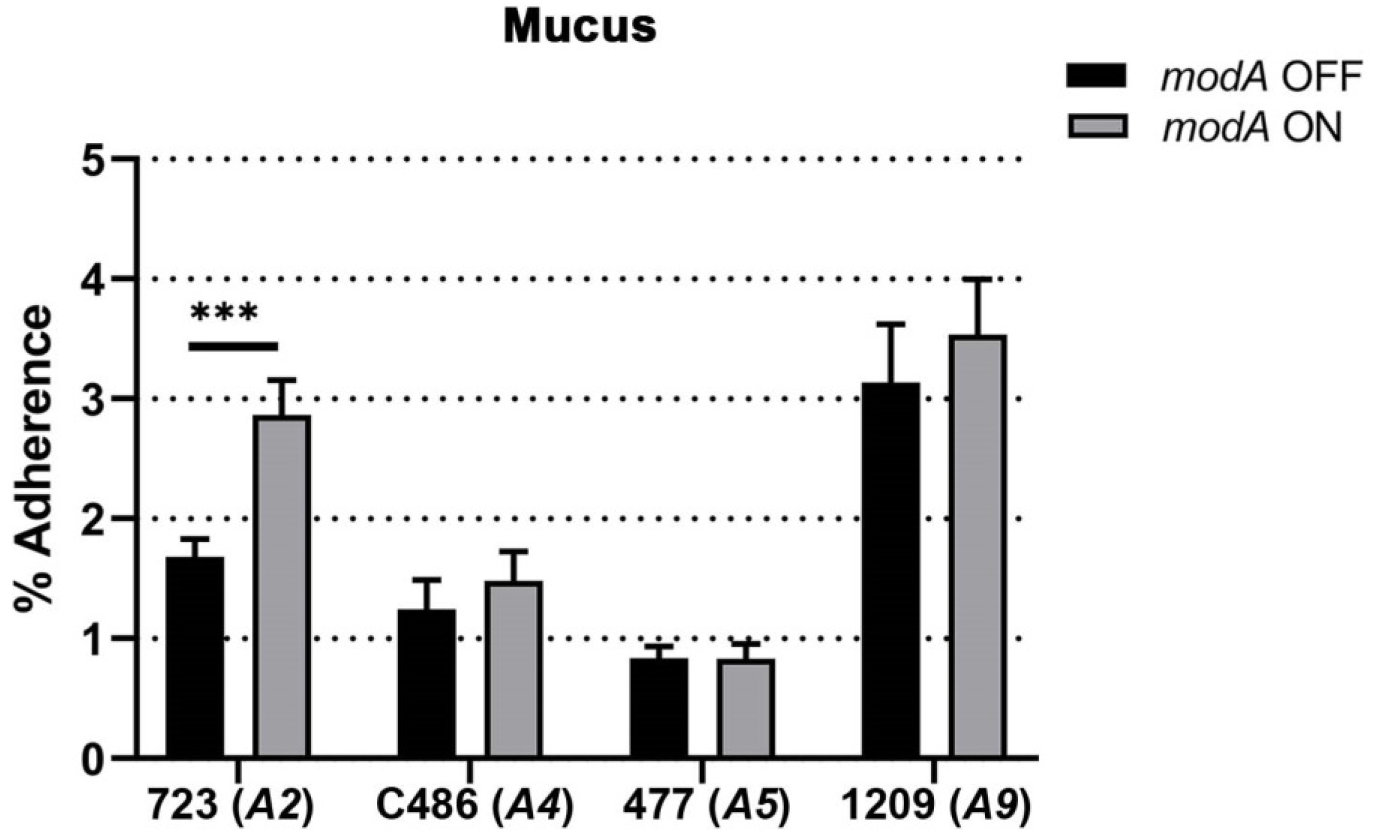
Adherence of NTHi *modA* locked variants to mucus. Data are shown as percentage of adherent bacteria relative to inoculum after 1 hour. The *modA2* ON variant adhered significantly better than the *modA2* OFF variant. There was no significant difference between the variants of *modA4, modA5* or *modA9*. *** *p* <0.001, Student’s t test.

### Adherence of NTHi to polarized and monolayer human bronchial-tracheal epithelial cells is not regulated by the ModA phasevarion

To assess the adherence of NTHi strains to pseudostratified respiratory tract epithelium, mucus was rinsed from the apical surface of polarized nhPBTEs and individual *modA* variants were allowed to adhere to the polarized cells for 1 hour. There was no significant difference between the adherence of any *modA* variant pairs. However, a ModA-independent difference was observed between the strains tested, as strains C486 and 1209 appeared to adhere better to the surface of polarized nhPBTEs in comparison to strains 723 and 477 (Fig. 2A).

**Fig. 2.**
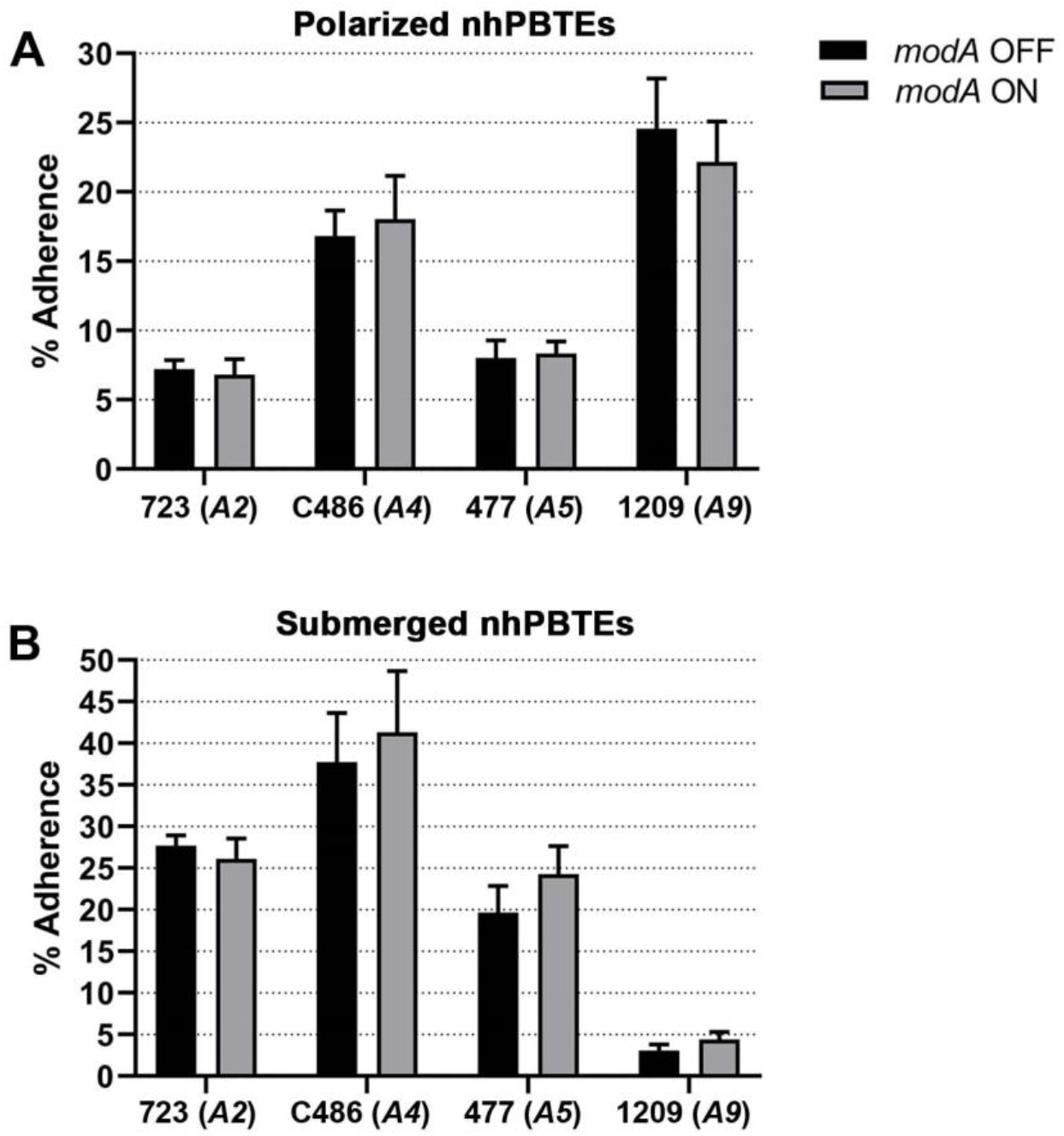
Adherence of NTHi *modA* locked variants to normal human primary bronchial-tracheal epithelial cells (nhPBTEs). Percentage of adherence to A) polarized nhPBTEs or B) submerged nhPBTEs after 1 hour was plotted. Percent adherence varied amongst strains; however, there was no significant difference between the *modA* variant pairs of any of the strains.

The complex pseudostratified structure of the polarized epithelium does not allow NTHi to access the basal epithelial cells underneath an intact airway epithelium (39). However, damage to the airway incurred during disease, or by environmental factors, can expose the basal epithelial layer to bacterial infections (41–43). Therefore, nhPBTEs were grown as monolayers submerged in culture medium to mimic the basal epithelial cells of the airway (44) and the adherence of the *modA* variants to these monolayers was assessed. Each strain adhered to a different extent, of which strain 1209 adhered the least (Fig. 2B), possibly due to general reduced expression of adhesins critical to adherence to these cells (33). This contrasted with the observation that strain 1209 adhered the greatest to polarized nhPBTEs, suggesting that different adhesins are required to adhere to submerged and polarized nhPBTEs. There was no ModA-dependent statistically significant difference observed between variant pairs of any of the strains (Fig. 2B). Therefore, ModA2, ModA4, ModA5 and ModA9 likely do not regulate the adherence of NTHi to pseudostratified or undifferentiated nhPBTEs.

### ModA2 and ModA9 regulate adherence of NTHi to middle ear epithelial cells

As the middle ear is a major site for NTHi infection, the middle ear epithelium is a common substrate for adherence by NTHi. Since the chinchilla is an established animal model to study the course of acute OM (45) and human middle ear cell lines are not readily available, chinchilla middle ear epithelial cells (CMEEs) were selected to study the adherence of NTHi to middle ear epithelium. CMEEs were cultured submerged in medium until confluent monolayers were formed and then adherence of the *modA* variants to CMEEs was assessed. The ability of all 4 strains to adhere to CMEEs (Fig. 3) was similar to that seen with submerged nhPBTEs (Fig. 2), as strain 1209 adhered the least in comparison to the other 3 strains. However, a ModA-dependent difference was observed between the variants of 723 and 1209. The *modA2* OFF variant adhered to CMEEs significantly more than the *modA2* ON variant (Fig. 3, p<0.0001), the reverse phenotype was demonstrated with adherence to mucus, whereas the *modA9* ON variant adhered significantly more than the *modA9* OFF variant (Fig. 3, p<0.05). The status of *modA* did not affect the adherence of strains C486 (*modA4*) or 477 (*modA5*) to CMEEs. Thus, ModA2 and ModA9 regulated the adherence of NTHi to middle ear epithelium in distinct ways.

**Fig. 3.**
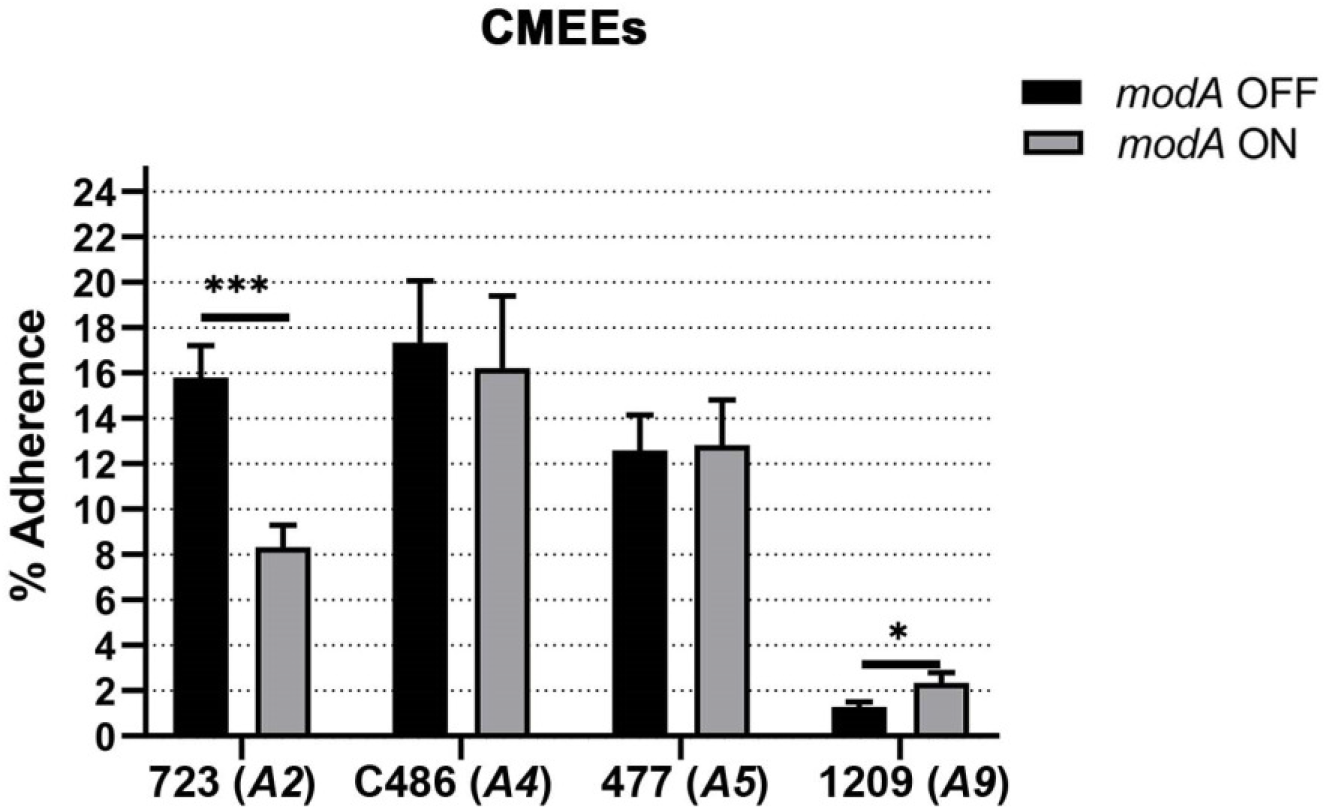
Adherence of NTHi *modA2* locked variants to chinchilla middle ear epithelial cells (CMEEs). Data are shown as percentage of adherent bacteria relative to inoculum after 1 hour. The *modA2* OFF variant adhered significantly better than the *modA2* ON variant and the *modA9* ON variant adhered significantly better than the *modA9* OFF variant. * *p* < 0.05 and *** *p* < 0.001, Student’s t test.

### ModA2 and ModA9 regulate adherence of NTHi to vitronectin

Adherence of NTHi to extracellular matrix (ECM) components is known to be important for adherence to host epithelial cells and survival of the pathogen during disease (46–49). Therefore, adherence of the *modA* variants to the ECM components fibronectin, laminin and vitronectin was assessed. While the *modA* status did not affect the adherence of any of the *modA* variant pairs to fibronectin or laminin (Fig 4A and B), significant differences in adherence to vitronectin were observed between the *modA* variant pairs of strains 723 (*modA2*) and 1209 (*modA9*). The *modA2* OFF variant adhered significantly better to vitronectin than the *modA2* ON variant (Fig. 4C, p=0.02). Similarly, the *modA9* OFF variant adhered significantly more than the *modA9* ON variant (Fig. 4C, p=0.001) which is in contrast to what we observed when relative adherence to CMEEs was assessed (see Fig. 3). Noticeably, strain 1209 adhered the least to all 3 ECM components whereas strain C486 adhered the most (Fig. 4A, B and C). Overall, these results suggest that ModA2 and ModA9 regulate the adherence of NTHi to vitronectin, possibly via the same adhesin(s).

**Fig. 4.**
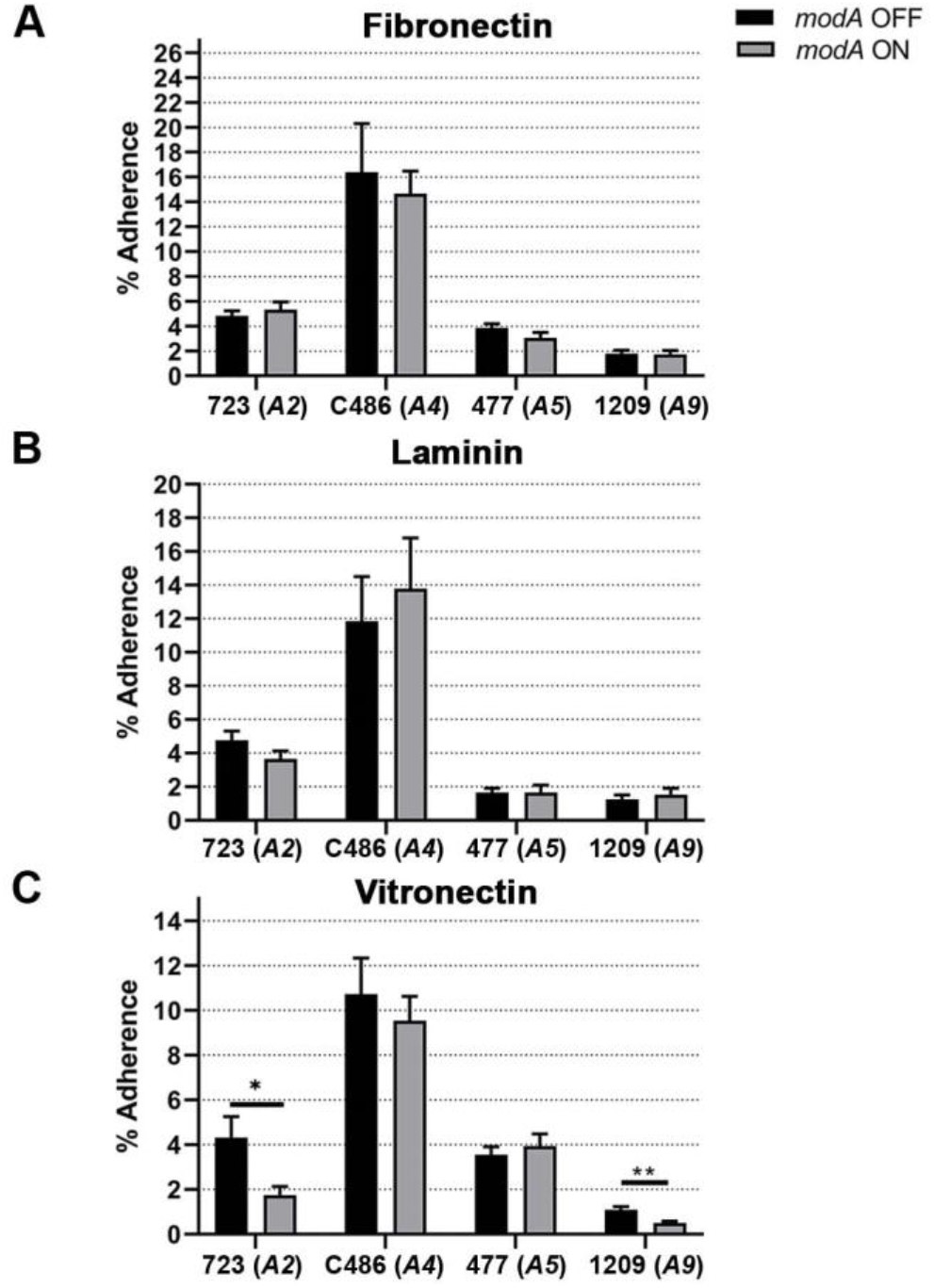
Adherence of NTHi *modA* locked variants to extracellular matrix components. Percentage of adherence to A) fibronectin, B) laminin and C) vitronectin after 1 hour was plotted. Strain-specific difference in adherence were observed for all ECM components. There were no ModA-specific differences in adherence to fibronectin or laminin for any of the strains. Adherence to vitronectin was significantly different between the *modA2* and *modA9* variant pairs. * *p* < 0.05 and ** *p* < 0.01, Student’s t-test.

### ModA2 regulates NTHi adherence dependent on adhesins PE and P4 in a substrate-specific manner

Of the various substrates tested, a clear correlation was observed in adherence of the NTHi strain 723 variants to vitronectin and CMEEs where the *modA2* OFF variant adhered significantly better than the *modA2* ON variant. Therefore, the contribution of NTHi adhesins required for adherence to vitronectin and epithelial cells in ModA2-dependent regulation was investigated.

The adhesin Protein E (PE) is known to mediate adherence of NTHi to vitronectin and thereby to respiratory epithelial cells (47, 49). The gene *ompE* that codes for PE was deleted from the genomes of the *modA2* locked variants. Deletion of *ompE* significantly reduced adherence of the *modA2* OFF variant to vitronectin (Fig. 5A, p<0.05) but did not change the adherence of the *modA2* ON variant (Fig. 5A, p=0.85). Since loss of *ompE* only affected the *modA2* OFF variant, PE may contribute to ModA2-dependent regulation of adherence to vitronectin. However, deletion of *ompE* from the *modA2* OFF variant did not completely reduce adherence to that of the *modA2* ON variant. Therefore, PE is likely not the only factor involved in ModA2-regulated adherence to vitronectin. A surface associated lipoprotein and adhesin, P4 (or outer membrane protein 4), encoded by the gene *hel* in NTHi, is also known to mediate adherence of NTHi to vitronectin and required for survival of NTHi in the middle ear (48). Deletion of *hel* significantly reduced the adherence of both variants to vitronectin (Fig. 5B). However, the percent adherence of the *modA2* OFF variant was reduced by 47% (Fig. 5B, 13.6% to 7.3%, p<0.001), whereas that of the *modA2* ON variant reduced by only 35% (Fig. 5B, 5.5% to 3.6%, p<0.05). This suggested that deletion of *hel* affected the *modA2* OFF variant to a greater extent than the *modA2* ON variant. Therefore, P4 may contribute to ModA2-dependent regulation of adherence of NTHi to vitronectin. The role of the adhesin Hap was also investigated in ModA2-regulated adherence to vitronectin and CMEEs by comparing the adherence of the *hap* deletion mutant variants with that of the wild-type variants. Deletion of *hap* did not affect adherence to vitronectin (Fig. 5C), which is not surprising as Hap is known to bind to the ECM components fibronectin, laminin and collagen IV but not vitronectin (46, 48).

**Fig. 5.**
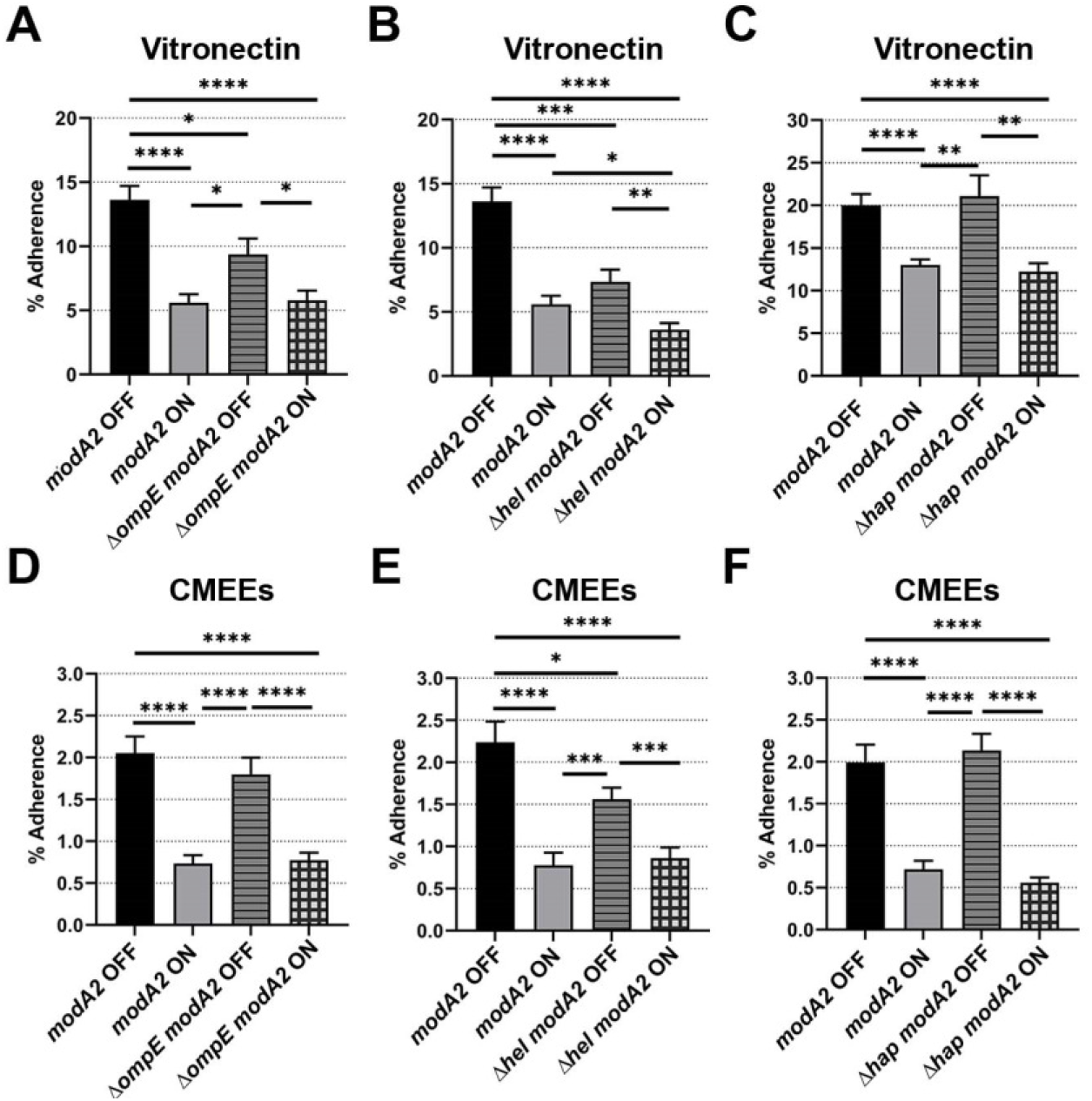
Role of adhesins PE (*ompE*), P4 (*hel*) and Hap (*hap*) in ModA2-dependent adherence of NTHi to vitronectin and CMEEs. Wild-type and adhesin mutants of the *modA2* OFF and the *modA2* ON variants of NTHi 723 were compared for adherence to vitronectin and CMEEs. Deletion of *ompE* significantly reduced the adherence of the *modA2* OFF variant to vitronectin but did not affect the *modA2* ON variant (A). Deletion of *hel* reduced the adherence of the *modA2* OFF variant to vitronectin more significantly than that of the *modA2* ON variant (B). Deletion of *ompE* did not affect the adherence of either variants to CMEEs (D), whereas deletion of *hel* significantly reduced the adherence of the *modA2* OFF variant and not the *modA2* ON variant (E). Deletion of *hap* did not affect the adherence of the *modA2* variants to vitronectin (C) and CMEEs (F). * *p-*value < 0.05, ** *p-*value < 0.01, *** *p-*value < 0.001 and **** *p-* value < 0.0001, Student’s t test.

Next, adherence of the adhesin mutant variants to CMEEs was assessed and compared with that of the wild-type variants. Interestingly, there was no effect of *ompE* deletion on the adherence of either variant to CMEEs (Fig. 5D). Therefore, PE may not be essential for adherence of strain 723 to CMEEs. In contrast, deletion of *hel* significantly reduced adherence of the *modA2* OFF variant to CMEEs but did not affect the *modA2* ON variant (Fig. 5E), suggesting that P4 may contribute to ModA2-dependent regulation of adherence to CMEEs. There was no significant difference between adherence of the *hap* mutant variants and the wild-type variants to CMEEs (Fig. 5F). Therefore, Hap may not contribute to the ModA2-dependent regulation of adherence to CMEEs. Taken together, both PE and P4 are likely involved in the ModA2-dependant regulation of adherence to vitronectin. Additionally, P4 may contribute to the ModA2-dependent regulation of adherence to CMEEs.

## Discussion

Adherence of NTHi to the host airway is critical for colonization as a commensal as well as for pathogenesis. Since the ModA phasevarion regulates the expression of surface associated proteins, including adhesins (33), and affects various aspects of pathogenesis of NTHi (33–36), this study aimed to address the role of ModA phasevarions in the adherence of NTHi to host airway components. The *modA* alleles *modA2, modA4, modA5, modA9*, and *modA10* are the most prevalent alleles in clinical isolates of NTHi collected from the nasopharynx of healthy individuals, and middle ears of OM patients, whereas the *modA2, modA4*, and *modA5* alleles are highly prevalent in the lungs of COPD patients (32, 33). Since the role of ModA10 is already reported in the adherence of NTHi to respiratory epithelial cells (50), the ModA2, ModA4, ModA5 and ModA9 phasevarions were selected for this study. The NTHi clinical strains 723, C486, 477 and 1209 were used to represent the ModA2, ModA4, ModA5 and ModA9 phasevarions, respectively. Variants of each of the 4 strains with the *modA* status locked to either OFF or ON allowed for assessment of the direct effect of each specific *modA* status. The adherence of the variant pairs of all 4 strains to different cellular and non-cellular substrates that are commonly encountered by NTHi within the airway was assessed. Epithelial cells from the middle ear of chinchillas (CMEEs) and normal human bronchial tracheal epithelial cells (nhPBTEs) were used as cellular models. Additionally, mucus and ECM components were used to represent non-cellular substrates of the respiratory tract. Adherence of all 4 strains varied based on the substrate either due to the inherent properties of the strains or the status of *modA*. This indicated the presence of a strict mode of regulation of adherence of NTHi that may favor a specific phenotypic variant to adapt better at different sites within the host airway.

This study demonstrated that the status of *modA2*, the most prevalent *modA* allele (33), significantly affected the adherence of NTHi to diverse respiratory tract substrates. The *modA2* ON variant of strain 723 adhered better to mucus, whereas the *modA2* OFF variant adhered better to CMEEs and vitronectin. However, ModA9 phasevarion regulated adherence of strain 1209 to CMEEs and vitronectin in an opposite manner. Strain 1209 adhered more to mucus and polarized epithelial cells than to submerged epithelial cells and ECM components, irrespective of *modA9* status. This could be due to the lack of expression of major adhesins like Hia and HMW in strain 1209 (33) that may be required to bind to cognate receptors on submerged epithelial cells. Moreover, pseudostratified epithelial cells secrete mucus unlike the submerged epithelial cells (44) and therefore can be adhered to better by strain 1209. Hence, cellular composition of the airway epithelium can dictate the ability of NTHi to adhere and cause disease. For example, strain 1209 may adhere more to the nasopharynx than to the middle ear epithelium due to the abundance of mucus producing goblet cells in the nasopharynx (51), whereas the reverse may occur in case of strain 723. Adherence of NTHi to submerged nhPBTEs also differed depending on *modA4, modA5* and *modA9* status but these differences were not statistically significant. Further, the NTHi *modA10* OFF variant has been reported to adhere better to human middle ear epithelial cells and bronchial epithelial cells compared to the *modA10* ON variant (50). Therefore, different ModA phasevarions affect the adherence of NTHi to different respiratory substrates in a unique manner, suggesting tissue- and niche-specific advantages conferred by the particular genes regulated by each phasevarion.

NTHi frequently adhere to mucus and ciliated epithelium within the nasopharynx (19). Although ModA2 regulated adherence to mucus, none of the ModA phasevarions affected adherence to pseudostratified epithelium. Since the ModA2, ModA4 and ModA9 phasevarions are reported to be prevalent in nasopharyngeal isolates recovered from healthy individuals (33), they may play a role in asymptomatic colonization of the nasopharynx. Although adherence of NTHi to ciliated epithelium was not regulated by these phasevarions, they may nonetheless affect colonization by alternate mechanisms. The movement of NTHi from the nasopharynx to the middle ear occurs via adherence to mucus within the Eustachian tube lumen when the upper airway is compromised by viral infection (19). Thus, multiple ModA phasevarions may play a role in adaptation of NTHi in the nasopharynx during environmental stress conditions such as viral infections. Such adaptation may further enable NTHi to reach different sites within the airway, leading to disease. As the *modA2* ON variant adhered to mucus better than the *modA2* OFF variant, the *modA2* ON status may be more advantageous for NTHi during initial colonization as well as ascension from the nasopharynx to the middle ear.

The ECM component vitronectin is reported to be detected in different parts of the airway (52–55). Since the *modA2* OFF variant adhered better than the *modA2* ON variant to vitronectin as well as CMEEs, ModA2 may regulate adhesins involved in the adherence to these two substrates. Although PE and P4 are both known to mediate adherence to vitronectin (48), each of these adhesins played a different role in the ModA2-dependent regulation of adherence. While ModA2-regulated adherence to vitronectin depended on PE and partially on P4, only P4 contributed to the ModA-regulated adherence to CMEEs. P4 is also known to play role in the virulence of NTHi in the middle ear (48). Hence, P4 may participate in ModA2-dependent regulation of adherence of NTHi to vitronectin as well as middle ear epithelium. On the other hand, PE may only contribute to the ModA2-dependent regulation of adherence to vitronectin. Therefore, ModA2 seems to affect adherence differently based on the type of host substrates. Interestingly, the transcription of *ompE* and *hel* has been found to be unaffected by the status of *modA2* (33), suggesting a role of ModA2 in the presentation or accessibility of these adhesins on the surface of NTHi. PE and P4 are also known to bind to fibronectin and laminin (48). However, ModA2 expression did not alter adherence of NTHi to fibronectin or laminin, indicating that the ModA2-regulated adherence to CMEEs may function independently of these 2 ECM components.

Nevertheless, adherence to vitronectin may not be the only mechanism NTHi use to adhere to CMEEs because ModA9 regulated the adherence to CMEEs and vitronectin with opposite trends. Since *modA9* is prevalent in OM isolates and not in COPD isolates (32, 33), the regulation of adherence to vitronectin by ModA9 may be important in the pathogenesis of NTHi during OM.

Vitronectin is also known to contribute to the repair of damaged airway epithelium during respiratory diseases (53–55). Studies have shown that certain clinical isolates of NTHi can associate better with damaged respiratory epithelium (56, 57). Therefore, the regulation of adherence to vitronectin by ModA2 could be important for adaptation of NTHi to the damaged or inflamed airway during diseases like COPD and CF. Although PE is known to be required for adherence of NTHi to bronchial epithelial cells via binding to vitronectin (49), the status of *modA2* did not affect adherence to submerged or polarized nhPBTEs. This could be due to the involvement of other bacterial and host factors, yet to be identified effects of ModA2 phasevarion, or limitations of the air-liquid interface (ALI) culture. For example, the Type IV pilus (Tfp) binds to the host receptor ICAM-1 (21) and high molecular weight (HMW) protein binds to various host proteoglycans (58). The HMW protein is also reported to be expressed more in the *modA2* ON variant than the *modA2* OFF variant (33). Therefore, it is possible that higher expression of adhesins other than PE may compensate for reduced adherence of the *modA2* ON variant to vitronectin on the surface of nhPBTEs.

Since *modA2* is the most prevalent *modA* allele found in clinical isolates of NTHi obtained from OM patients (33), the ModA2-dependent regulation of adherence to mucus, vitronectin and middle ear epithelial cells likely contributes to NTHi pathogenesis during OM. Greater adherence to mucus may provide the *modA2* ON variant with an advantage in ascending the Eustachian tube into the middle ear (19). We have previously reported that the *modA2* ON status is selected for within middle ear fluids during a chinchilla model of experimental otitis media, and that a shift from *modA2* OFF to ON status occurs within the middle ear that leads to a more severe disease pathology than when the middle ear is initially challenged with a predominantly *modA2* ON population (33, 36). Overall, these data suggest that NTHi may enter the middle ear predominantly in the *modA2* ON status following ascension from the nasopharynx. However, once in the middle ear, NTHi encounter niche-specific microenvironmental conditions and host defenses. Regulation by ModA2 results in a population that consists of bacteria in both the *modA2* ON status and the *modA2* OFF status at varied proportions. Each of the sub-populations express a unique set of virulence factors and is thus primed to survive different stressors. We have shown herein that the *modA2* OFF variant is better able to adhere to the middle ear epithelium and vitronectin, which may cause this variant to attach to the epithelial surface more than the *modA2* ON variant. The *modA2* ON variant may instead remain predominantly in the effusion. The presence of NTHi within the middle ear will also lead to infiltration of immune cells, like neutrophils and macrophages (37). The ability of the *modA2* ON variant to resist killing by macrophages could further contribute to the selection of the *modA2* ON population in the middle ear fluids (37). On the other hand, the *modA2* OFF variant is known to better resist neutrophil-mediated killing and oxidative stress (34, 37) and form more stable biofilms under environmental conditions found during OM (35), all factors that may result in better survival of the *modA2* OFF variant at the mucosal surface and within the mucosal biomass that is predominantly composed of neutrophils and extracellular traps (NETs). Future studies to determine the localized variant selection within the middle ear during disease are of interest, however, are technically challenging due to the robust immune response within the mucosal biomass. Overall, the compartmentalization of adherence by the *modA2* variants may assist in niche-specific adaptation of NTHi during the disease course.

In conclusion, this study established that the clinically prevalent ModA phasevarions regulate adherence of NTHi to specific host airway substrates. These findings are important because adhesins are well known vaccine candidates against NTHi (59–63). Adhesins have also been useful in the diagnosis of NTHi-induced respiratory infections (64) and can be targeted for eradication of adherent NTHi population from the site of disease (65). Since each *modA* variant has a certain advantage over the other, the switch from one *modA* status to another may enable NTHi to evade recognition and clearance by the host immune response or therapeutic agents. Therefore, understanding the regulation of adherence by the ModA phasevarion at the site of colonization and disease is necessary to increase the potential use of adhesins as vaccine candidates, diagnostic tools, and therapeutic agents against NTHi.

## Materials and methods

### Bacterial strains and growth conditions

NTHi strains 723, 477 and 1209 were received from the Finnish Otitis Media study group (66) and strain C486 was isolated from a child with otitis media (67). The *modA* locked variants of each of these strains were constructed as described previously (Brockman et al., 2017), so that these strains were unable to switch the status of *modA*. NTHi strains were cultured at 37°C and 5% CO_2_ on chocolate agar or in brain heart infusion (BHI) broth supplemented with hemin (2 μg/mL) and β-NAD (2 μg/mL). All strains used are listed in Table 1.

### Generation of mutants

DNA fragments containing a kanamycin resistance gene flanked by sequences homologous to the sequences flanking the target genes were designed and then synthesized (Integrated DNA Technologies). Each fragment was ligated into a pJET1.2 blunt end cloning vector at the EcoRV restriction site using a CloneJET PCR cloning kit (Thermo Scientific). *E. coli* DH10B competent cells were transformed with the ligation products and transformant colonies were selected on LB agar plates containing ampicillin. Plasmids isolated from these transformant colonies were used as templates for amplifying the inserts using the pJET1.2 forward and reverse sequencing primers (Table 2). NTHi 723 *modA2* locked OFF and *modA2* locked ON variant strains were transformed with the amplified inserts using the M-IV method (68). NTHi transformants were selected on chocolate agar plates supplemented with kanamycin. The mutants were confirmed by sequencing as well as by PCR using kanamycin resistance cassette internal primers and primers for flanking sequences of the target gene. All primers used for cloning and PCR are listed in Table 2.

**Table 2:**
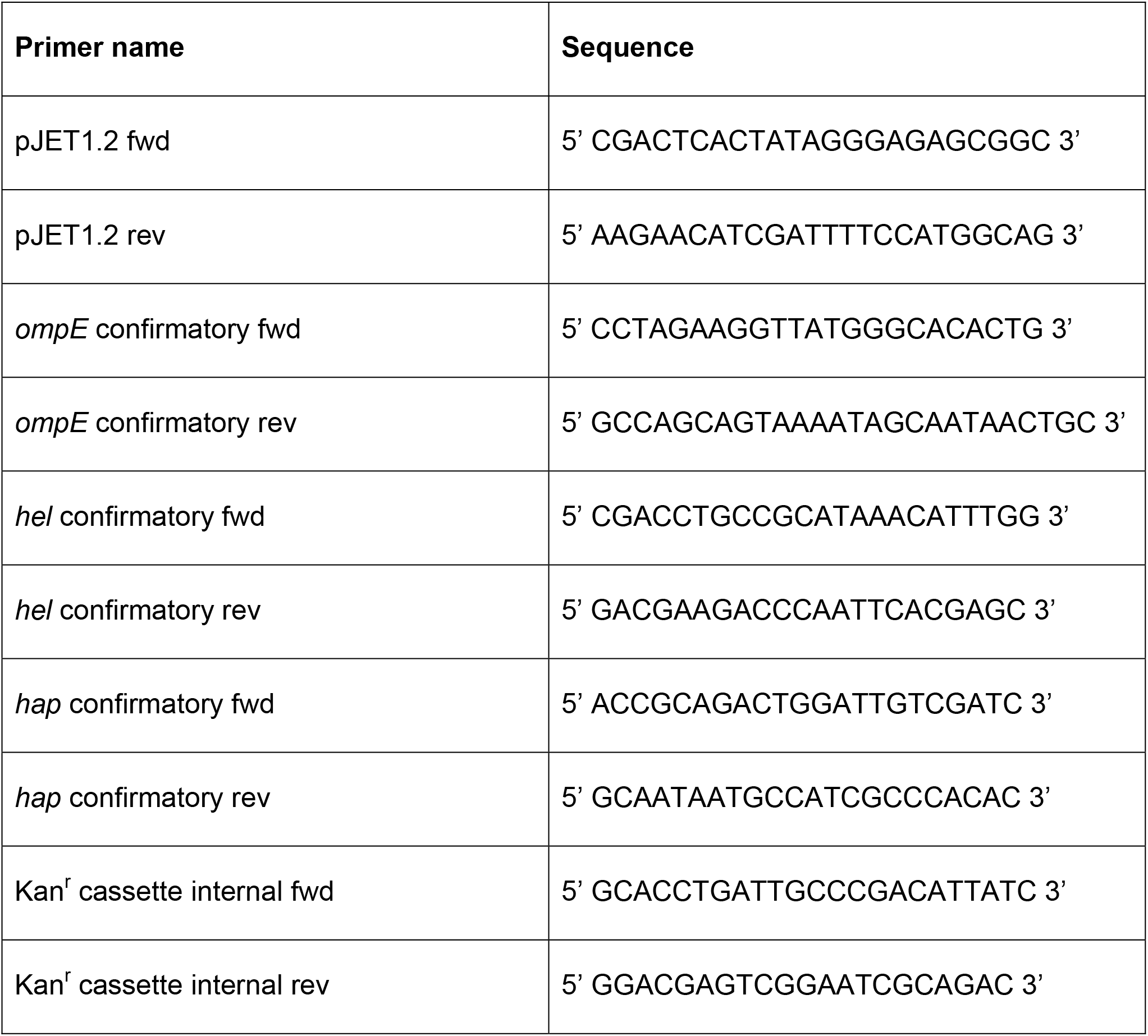
Primers.

### Isolation and culturing of CMEEs

Chinchilla middle ear tissues were aseptically harvested from naive animals and cultured in explant medium (69) containing DMEM (Mediatech), Ham’s F12 (Mediatech), Glutamine (Mediatech), Hydrocortisone (STEMCELL TECHNOLOGIES), Isoproterenol (Sigma) and fetal bovine serum (FBS) (Mediatech). Epithelial cells (CMEEs) generated from the tissues were maintained in explant medium containing epidermal growth factor (EGF) (Sigma).

### Adherence assays with CMEEs

CMEEs were seeded into wells of 96-well plates (Costar, flat bottom) and maintained in culture medium until the formation of a tightly packed monolayer. Bacterial inoculum was prepared from log phase cultures of NTHi variant pairs and added to wells containing the CMEEs at an MOI of 100. The cells were incubated at 37°C and 5% CO_2_ for 1 hour to allow the bacteria to adhere. The medium was removed and the CMEEs were washed 3 times with 1X DPBS to remove non-adherent bacteria. 10X TrypLE (Gibco) was used to detach the adherent bacteria which were then collected in 1X DPBS, serially diluted and plated on chocolate agar. Percent adherence was determined from the CFU values of the adherent bacteria and the inoculum.

### Adherence assays with submerged nhPBTEs

Normal human primary bronchial-tracheal epithelial cells (nhPBTEs) obtained from healthy human lungs (ATCC PCS-300-010) were seeded into the wells of 96-well plates (Costar, flat bottom) and maintained in Pneuma-Cult Expansion medium (PC-Ex) (STEMCELL TECHNOLOGIES) till confluency. Bacterial inoculum was prepared from log phase cultures of NTHi and added to wells containing the nhPBTEs at an MOI of 100. The cells were incubated for 1 hour at 37°C and 5% CO_2_. Medium was removed and cells were washed 3 times with 1X DPBS to remove non-adherent bacteria. 10X TrypLE (Gibco) was used to detach the adherent bacteria which were then collected in 1X DPBS and plated on chocolate agar. Percent adherence was determined as described above.

### Adherence assays with polarized nhPBTEs

nhPBTEs were seeded in 6.5 mm Transwells (Corning Transwells) and maintained in PC-Ex expansion medium. Upon reaching confluency, media was removed from the apical surface and cells were fed basolaterally with Pneuma-Cult ALI (Air Liquid Interface) differentiation medium (STEMCELL TECHNOLOGIES) for 5-8 weeks to allow differentiation of cells at the air-liquid interface. For the assay, the apical surface was washed with 1X DPBS to remove mucus produced by these cells. Bacterial inoculum was prepared from a log phase culture of NTHi, added to the cells at an MOI of 100 and incubated at 37°C, and 5% CO_2_ for 1 hour. After the incubation period, the supernatant was removed and the apical surface was washed 3 times with 1X DPBS to remove non-adherent bacteria. 10X TrypLE (Gibco) was added to the apical surface and 1X TrypLE was added to the basolateral surface to dissociate the adherent bacteria. Samples were collected in 1X DPBS and plated on chocolate agar. Percent adherence was determined as described above.

### Mucus collection and quantification

nhPBTEs were differentiated into polarized cells at the air-liquid interface as described above and cultured until mucus production was observed. The apical surface was incubated with 1X DPBS at 37°C for 15 min and the mucus was collected by pipetting. Mucus was quantified using a Qubit protein assay kit and then added into the wells of Nunc MaxiSorp flat bottom 96-well plates at a concentration of 10 μg/well and incubated overnight at 37°C.

### Adherence assays with mucus

Mucus was collected, quantified, and used to coat Nunc MaxiSorp flat bottom 96-well plates as described above. Prior to the assay, mucus-coated wells were washed 4 times with 1X DPBS to remove excess mucus. Bacterial inoculum was prepared from log phase cultures of NTHi in 1X DPBS and added at a density of 5×10^6^ CFU/well. After 1 hour of incubation at 37°C and 5% CO_2_, the supernatant was removed and wells were washed 4 times with 1X DPBS to remove non-adherent bacteria. 1X DPBS (100 uL) was then added to each well and the adherent bacteria were dislodged and collected by vigorous pipetting and scraping of the wells. Dilutions of the collected samples were plated on chocolate agar. Percent adherence was determined using CFU values as described above.

### Adherence assays with ECM components

Flat bottom 96-well tissue culture treated plates (Costar) were coated with fibronectin (Sigma-Aldrich), laminin (Sigma-Aldrich) or vitronectin (Sigma-Aldrich) according to manufacturer protocols. Briefly, working solutions of vitronectin (1.5 ug/ml) and laminin (6 ug/ml) were prepared in 1X DPBS, whereas fibronectin was reconstituted in water (15 ug/ml). 100 ul of the working stock of vitronectin was added per well of 96 well plate and incubated at 37°C for 2 hours followed by overnight incubation at 4°C. Alternatively, 100 ul of the working stocks of laminin and fibronectin were added per well on the day of the assay, removed immediately and allowed to dry. The coated wells were washed twice with 1X DPBS just prior to the assay. Bacterial inoculum was prepared from log phase cultures of NTHi and added to coated wells at a density of 5×10^6^ CFU/well. After incubation at 37°C and 5% CO_2_ for 1 hour, the supernatant was removed, and wells were washed 4 times with 1X DPBS to remove any non-adherent bacteria. Adherent bacteria were collected in 100 ul 1X DPBS with vigorous pipetting and scraping of the wells. Dilutions of the collected sample were serially diluted and plated on chocolate agar. Percent adherence was determined as described above.

### Statistical analysis

Statistical significance was assessed by Student’s unpaired t-test using GraphPad Prism software, version 8.4.2. A *p-*value less than or equal to 0.05 was considered as statistically significant. Each experiment was carried out at least 3 times with a minimum of 3 biological replicates each time.

## Acknowledgements

This work was supported by the National Institute of Health (NIH) / National Institute of Deafness and other Communication Disorders (NIDCD) grants NIH/NIDCD R01DC015688 to L.O.B and M.P.J, and NIH/NIDCD R21DC016709 to K.L.B, and by the National Health and Medical Research Council Principal Research Fellowship 1138466 to M.P.J.

